# Whole genome bisulfite sequencing reveals a sparse, but robust pattern of DNA methylation in the *Dictyostelium discoideum* genome

**DOI:** 10.1101/166033

**Authors:** Jacob L. Steenwyk, James St. Denis, Jacqueline M. Dresch, Denis A. Larochelle, Robert A. Drewell

## Abstract

DNA methylation, the addition of a methyl (CH_3_) group to a cytosine residue, is an evolutionarily conserved epigenetic mark involved in a number of different biological functions in eukaryotes, including transcriptional regulation, chromatin structural organization, cellular differentiation and development. In the slime mold *Dictyostelium*, previous studies have shown the existence of a DNA methyltransferase (DNMA) belonging to the DNMT2 family, but the extent and function of 5-methyl-cytosine in the genome is unclear. Here we present the whole genome DNA methylation profile of *Dictyostelium discoideum* using deep coverage, replicate sequencing of bisulfite converted gDNA extracted from post-starvation cells. We find an overall very low level of DNA methylation, occurring at only 462 out of the ~7.5 million (0.006%) cytosines in the genome. Despite this sparse profile, significant methylation can be detected at 51 of these sites in replicate experiments, suggesting they are robust targets for DNA methylation. These 5-methyl-cytosines are associated with a broad range of protein-coding genes, tRNA-encoding genes and retrotransposable elements. Our data provides evidence of a minimal, but functional, methylome in *Dictyostelium*, thereby making *Dictyostelium* a candidate model organism to further investigate the evolutionary function of DNA methylation.

## Introduction

DNA methylation is a post-synthetic modification that typically occurs on cytosine residues in eukaryotic plants and animals [1]. Generally, DNA methylation is associated with transcriptional repression [2], but has been linked to more complex processes including cell differentiation [3], genomic imprinting and stability [4, 5], and X chromosome inactivation [6]. The majority of DNA methylation is carried out by the DNA methyltransferases DNMT1, DNMT3A and DNMT3B [7]. DNMT1 is responsible for maintaining a cell’s methylation profile post-replication by targeting hemi-methylated sites in the genome [8], while DNMT3A and DNMT3B methylate CpG dinucleotides *de novo*, thereby creating new epigenetic marks [9]. Although DNA methylation appears to be evolutionarily conserved across a large number of eukaryotes (Fig. 1), some organisms, such as *Dictyostelium discoideum* and *Drosophila melanogaster*, lack the major DNMTs while retaining the less characterized methyltransferase DNMT2 [8].

**Figure 1.**
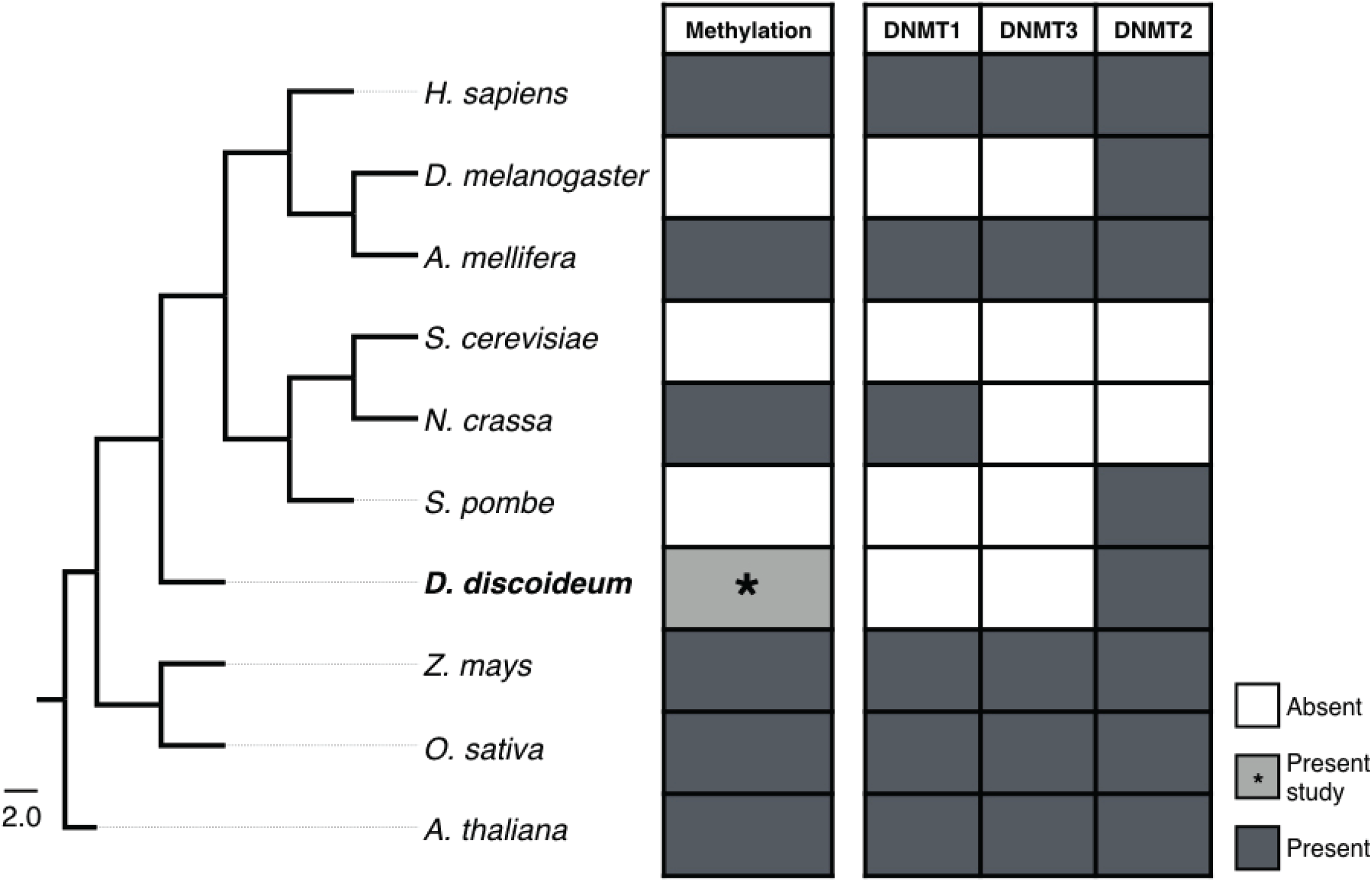
Evolution of DNA methyltransferases. The presence (dark boxes) or absence (white boxes) of biologically significant DNA methylation and the DNA methyltransferase (DNMT) enzymes across eukaryotes. Kingdom, Animalia: H. sapiens, Homo sapiens; *D. melanogaster*, *Drosophila melanogaster; A. mellifera, Apis mellifera. Kingdom, Fungi: S. cerevisiae*, *Saccharomyces cerevisiae*; *N. crassa, Neurospora crassa; S. pombe, Schizosaccharomyces pombe*. Kingdom, Protozoa: *D. discoideum, Dictyostelium discoideum*. Kingdom Plantae: *Z. mays, Zea mays* subsp. *mays; O. sativa, Oryza sativa; A. thaliana, Arabidopsis thaliana*. The tree was generated using the NCBI taxonomy common tree (https://www.ncbi.nlm.nih.gov/Taxonomy/CommonTree/wwwcmt.cgi).

DNMT2 was originally identified based on sequence conservation of essential catalytic motifs and exhibits structural similarity to other DNMTs [10, 11]. It is found as a single copy gene across the eukaryotic tree of life in protists, plants, fungi and animals (Fig. 1), suggesting an important functional role [11]. To date, DNMT2 has been shown to methylate tRNAs [12] and contribute to DNA methylation of retrotransposons in *D. melanogaster* [13], demonstrating a dual specificity for DNA and RNA substrates [14]. Despite these observations, characterizing the presence, relevance and efficacy of DNA methylation in organisms containing only a DNMT2 methyltransferase remains an active area of research [10, 11, 14-16].

*Dictyostelium discoideum*, a eukaryotic slime mold containing a *Dnmt2*-homolog *(dnmA/DDB0231095*) [17, 18], is among the organisms where the presence, extent and function of DNA methylation is debated. In the initial 1991 study, *D. discoideum* was reported to lack DNA methylation, as determined by whole genome methylation-sensitive restriction enzyme analysis and high performance liquid chromatography assays [19]. Despite this early report, the confirmed existence of a DNMT2 homolog, along with *D. discoideum’s* unique AT-rich genome and an under-representation of CpG dinucleotides relative to the GpC isomer, implied the possible presence of a DNA methylation system, as methylated CpGs are inherently chemically unstable and readily mutate to TpGs [17]. Accordingly, the investigation of DNA methylation in D. *discoideum* was revisited in 2006 using more advanced methodologies including antibodies to detect 5-methylcytosine in bulk genomic DNA across distinct developmental time points [20]. These studies detected overall low levels of DNA methylation in the genome and were confirmed using methylation-sensitive restriction digests targeted to retrotransposons and several other genes [20]. A functional role for the DNMA enzyme in silencing of retrotransposons via the asymmetric methylation of cytosine residues was confirmed using bisulfite sequencing of specific sites [18]. Furthermore, DNA methylation was demonstrated to increase through development (with the highest levels at 24 hours post-starvation) and knocking out dnmA revealed developmental defects and reduced DNA methylation [20]. More recently, detailed studies examining the activity of the *D. discoideum* DNMA enzyme demonstrated that specific tRNA molecules are the preferred target substrate for this methyltransferase, but other substrates potentially remain to be characterized [21]. These conflicting reports, coupled to the relatively few studies investigating DNA methylation in *D. discoideum*, therefore reflect the uncertainty regarding the status of the DNA methylation system in this species.

To determine if a functioning DNA methylation system is present in *D. discoideum*, we utilized whole genome bisulfite sequencing [22] to give deep coverage (~435x) of genomic DNA isolated from cells in an 18-24 hours developmental time window. Our results show that *D. discoideum* harbors a minimal, but replicable methylome. Overall, cytosine methylation is sparse in the *D. discoideum* genome, occurring at only 0.006% of all sites (462 out of the ~7.5 million cytosines in the genome), many of which demonstrate low levels of methylation. Despite this profile, significant methylation can be consistently detected at 51 of these sites in replicate experiments, suggesting they are robust targets for DNA methylation. These 5-methylcytosines are associated with a broad range of protein-coding genes, tRNA-encoding genes and retrotransposable elements. Taken together, our studies provide evidence of a minimal, but functional, methylome in *D. discoideum*, thereby making *D. discoideum* a candidate model organism to further investigate the enzymatic role of *Dnmt2* and the evolutionary function of DNA methylation.

## Results and Discussion

### Mapping, conversion and coverage rates

Sequencing of bisulfite-converted genomic DNA from *Dictyostelium discoideum* AX4 cells grown to between 18 and 24 hours of development, and spike-in control lambda bacteriophage DNA, generated 125 million reads after quality control (see Methods for details). This time window for the cells was chosen to ensure temporal heterogeneity of the gDNA sample and reflects the fact that previous studies have reported higher levels of DNA methylation in cells 24 hours post-starvation [20]. Of the sequencing reads, 72.53% mapped to unique regions of the *D. discoideum* genome, which equates to 286x coverage across the entire genome. More than 90% of the bases in the genome are covered by 100 or more reads (Fig. 2a and b). Over 80% of cytosines are covered by 75 or more reads (Fig. 2c), resulting in coverage of 98.816% of all cytosines in the genome (Table S1) with very similar profiles for CG, CHG and CHH sequence context (Fig. 2c). The bisulfite conversion rate in our experiment was 99.44%, measured by the C-to-U deamination rate in the unmethylated lambda genome, indicating that the false-negative rate was less than 0.6%.

**Figure 2.**
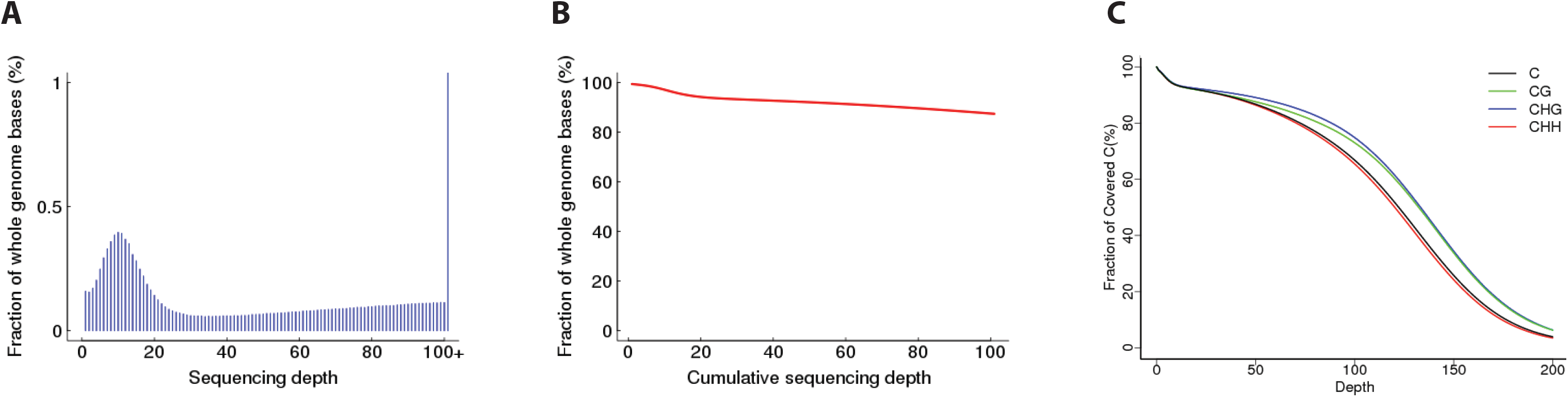
Sequencing depth and cumulative coverage of the Dictyostelium methylome. **(A)** Sequencing depth distribution across the entire genome. **(B)** Cumulative coverage across the entire genome, e.g. approximately 90% of sites are covered by 100 or more reads. **(C)** Percentage of cytosines that have a certain level of coverage, e.g. approximately 80% of sites have coverage of 75 reads. The profile for cytosines in different sequence contexts (CG, CHG and CHH, where H = non-G base) was similar.

### Overall genomic methylation profile

A very small proportion of cytosines in the *Dictyostelium* genome are methylated. The average overall level of methylation across the entire genome, calculated from the ratio of C reads to total reads (see Methods for details), is only 0.523%, with a similar profile across all chromosomes and mitochondrial DNA (Table 1). It should be noted that this may include a number of false positive C reads arising from a failure to convert in the bisulfite reaction and therefore is likely an overestimate of the global methylation level. Nonetheless, methylation is detected predominantly at CHH sequences (where H = non-G base), but is also found at CG and CHG sequences (Table 2). Of the approximately 7.5 million cytosines in the genome, only 462 (0.006% of all sites) are significantly methylated, of which only 27 are CpG sites (Table 2). This methylome profile mirrors previous studies that have reported global methylation levels in the *Dictyostelium* genome of 0% [19], 0.14% [20], and 0.20% [18] using different methodologies, indicating that DNA methylation is very rare in this species. These results are strikingly different from the methylation profile observed in other eukaryotes. In mammals, 60-90% of CpGs can be methylated [23], while in hymenoptera insects 0.51-0.67% of CpGs are methylated [24, 25]. The sparse 0.006% of methyl-CpGs detected in *Dictyostelium* is therefore at least two orders of magnitude lower than the level found in other eukaryotic organisms. Among the individual sites that demonstrate a significant level of methylation in our study, the vast majority are methylated in less than 10% of reads, but cytosines with up to a 100% methylation level are detected (Fig. 3).

**Table 1.** Methylation level by chromosome. The values represent the average methylation level at cytosine bases on each chromosome and the mitochondrial DNA (chrMT). The percentage value was determined by the ratio of the number of reads supporting methylation to the number of reads covering a particular cytosine site (see methods for details). The profile for cytosines in different sequence contexts (CG, CHG and CHH, where H = non-G base) is shown. It should be noted that the calculated bisulfite conversion rate in the experiment was 99.44%.

| Chromosome | C | CG | CHG | CHH |
| --- | --- | --- | --- | --- |
| chr1 | 0.519 | 0.438 | 0.481 | 0.531 |
| chr2 | 0.525 | 0.435 | 0.484 | 0.537 |
| chr3 | 0.524 | 0.439 | 0.484 | 0.536 |
| chr4 | 0.523 | 0.434 | 0.487 | 0.535 |
| chr5 | 0.521 | 0.432 | 0.484 | 0.532 |
| chr6 | 0.521 | 0.435 | 0.481 | 0.533 |
| chrMT | 0.528 | 0.46 | 0.499 | 0.546 |
| Whole Genome | 0.523 | 0.442 | 0.483 | 0.537 |

**Table 2.** Methylated cytosines in CG, CHG and CHH genomic sequence context. Methylation occurs predominantly at CHH sites, but is also detectable at CG and CHG sites. Of the approximately 7.5 million cytosines in the genome, only 462 (0.006% of all sites) are significantly methylated.

|  | mCG | mCHG | mCHH | mC |
| --- | --- | --- | --- | --- |
| **mC reads** | 180 | 161 | 3,431 | 3,772 |
| **proportion (%)** | 4.772 | 4.268 | 90.960 | 100 |
| **mC sites** | 27 | 19 | 416 | 462 |
| **proportion (%)** | 5.844 | 4.113 | 90.043 | 100 |
| **total C number** | 442,821 | 733,151 | 6,317,717 | 7,493,689 |
| **proportion mC (%)** | 0.006 | 0.003 | 0.007 | 0.006 |

**Figure 3.**
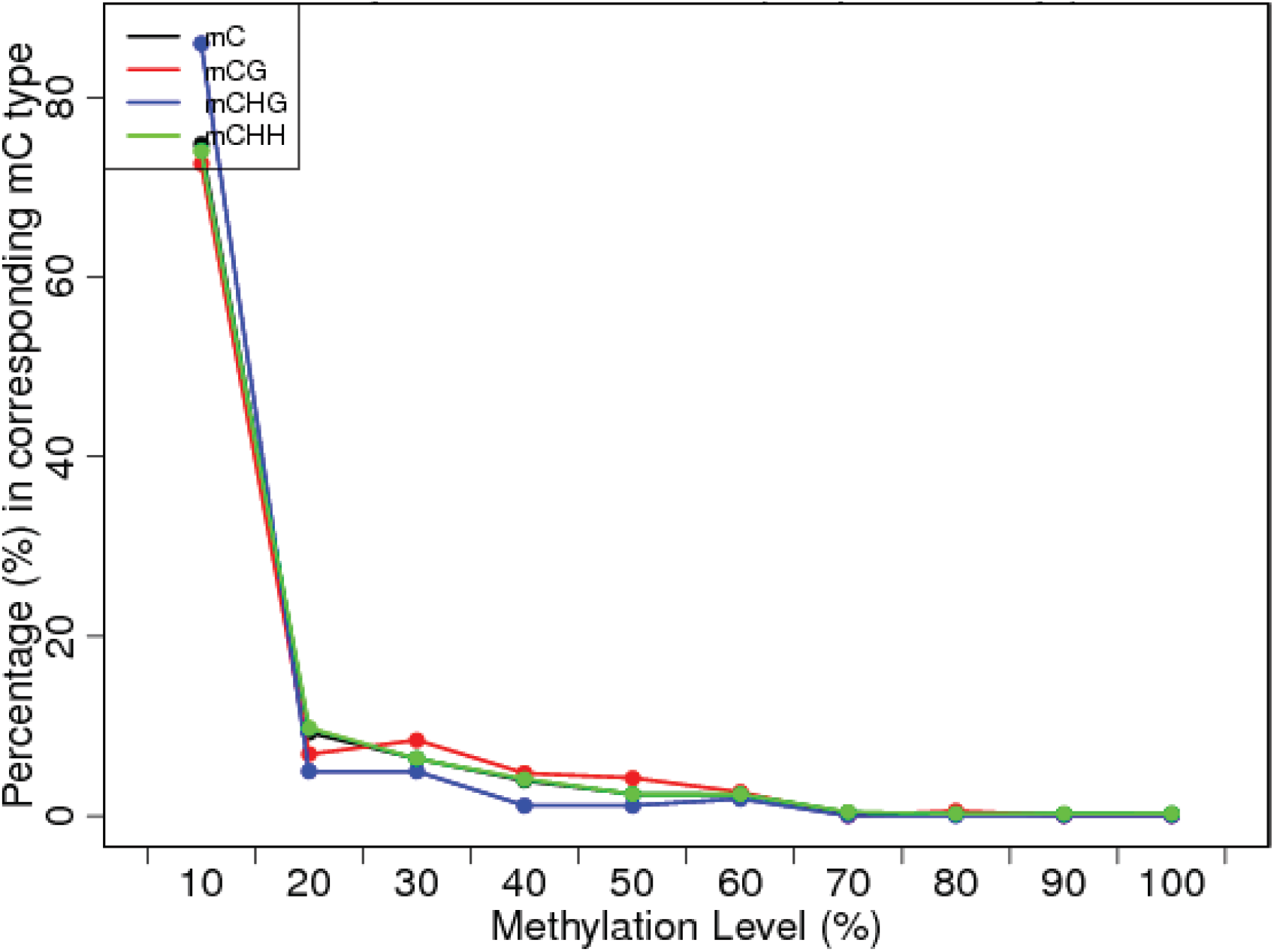
Overall methylation level at methyl-cytosines. The methylation level at the 462 individual mCs is calculated and organized in 10% bins (i.e. 0-10%, 91-100% etc.). Only mCs covered by at least four and no more than 1000 reads are used in this calculation. The vast majority of mCs, irrespective of sequence context, have less than 10% methylation level.

### Methylation levels at individual mCs

In order to further investigate the methylation level at the 462 individual cytosine residues we analyzed their distribution across the genome. All chromosomes and the mitochondrial DNA show a wide range of methylation levels for mCs, with mCHH sites carrying the broadest range (Fig. 4a). In parallel, we plotted the total read number for each of these mCs (Fig. 4b) against methylation level. Strikingly, there is a clear inverse relationship between read depth and methylation level at the mCs, with sites showing higher levels of methylation having lower read depth and vice-versa (Fig. 4c and d). This indicates that the high level of methylation detected at some of these mCs may simply reflect the overall low number of reads at these sites.

**Figure 4.**
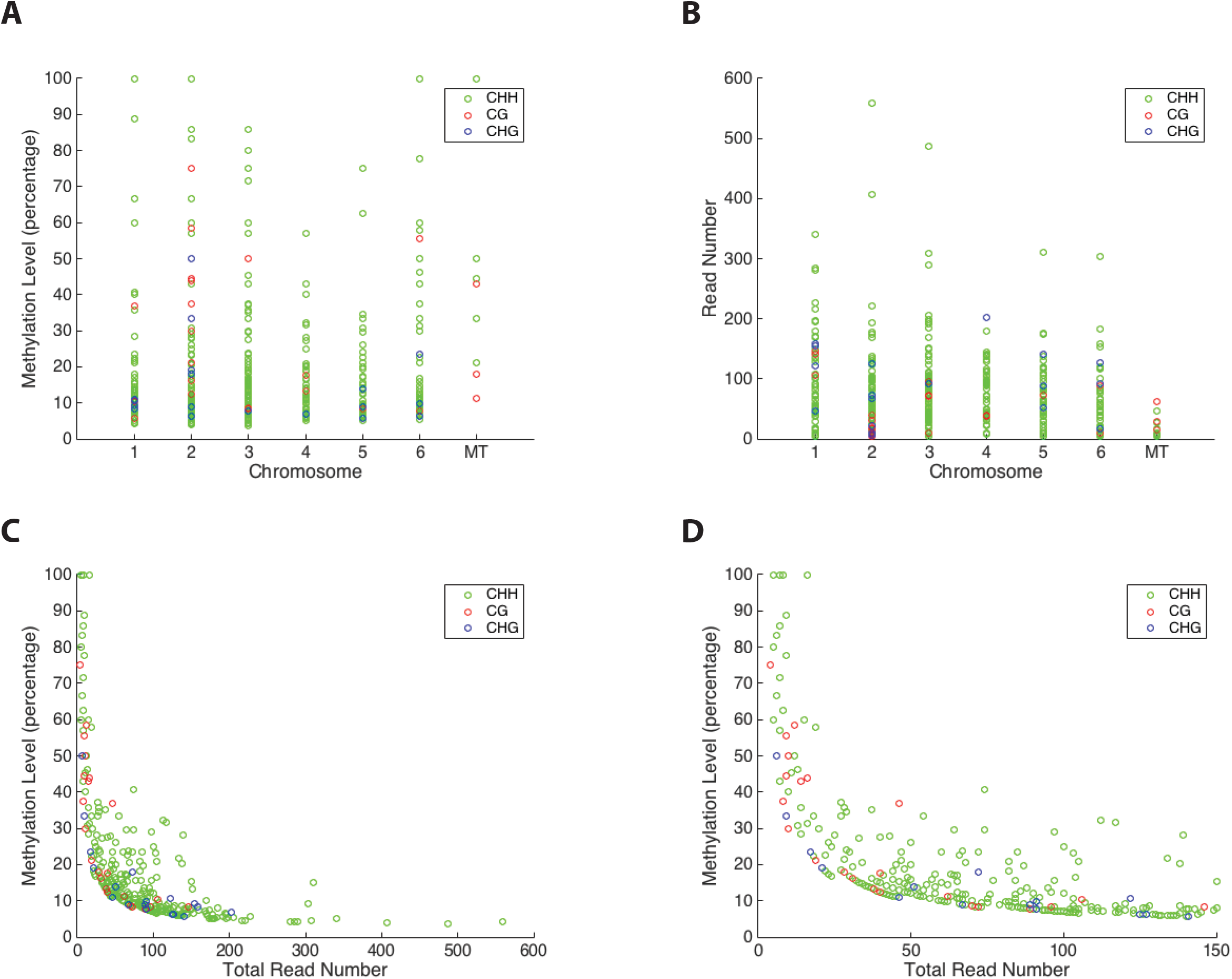
Methylation level and read depth at individual mCs. **(A)** The methylation level at the 462 individual mCs is depicted for each of the six genomic chromosomes and the mitochondrial (MT) DNA. Only mCs covered by at least four and no more than 1000 reads are shown. **(B)** The total read number for the 462 individual methylated sites. The vast majority of mCs, irrespective of sequence context, have less than 100 total reads. **(C)** and **(D)** The methylation level (%) plotted against the total read number for the 462 individual methylated sites demonstrates a clear inverse relationship between read depth and methylation level at the mCs, with sites showing higher levels of methylation having lower read depth and vice-versa. All the methylated sites in the genome with a methylation higher than 40% have fewer than 25 total reads.

### Comparative analysis between replicate methylomes

In an effort to examine the *Dictyostelium* methylome profile in more detail, we performed a replicate of the entire experiment. The overall results for the second-round of whole genome methylation analysis were comparable to the first-round results (see Table 3 for comparative summary. The same general characteristics for the two replicate methylomes were observed, including a very low level of methylation with just 462 and 459 significantly methylated cytosines, respectively, in the entire genome (Table 3). Surprisingly, 51 of the mCs were shared between the two replicates, including 5 mCGs, 2 mCHGs, and 44 mCHHs (Table 4), representing 11.04% of all the mCs. At the vast majority of these 51 individual mC sites, the level of methylation is above 10% and the total number of reads is greater than 10 in both replicates (Table 4). These 51 robust mCs are widely dispersed across the *Dictyostelium* genome, with numerous mCs found on each of the six chromosomes and in the mitochondrial DNA (Fig. 5). An additional feature found amongst the mCHH sites is that 20 of the 44 methylated cytosines are found in close proximity pairs (shown in gray in Table 4, Table S2).

**Table 3.** Comparison between the two replicate methylomes. Key metrics for the replicate methylomes. The overall profile is similar in both experiments.

|  | First-round | Second-round |
| --- | --- | --- |
| Clean reads | 125,357,640 | 73,253,322 |
| Mapping rate (%) | 72.53 | 67.96 |
| Bisulfite conversion rate (%) | 99.44 | 99.50 |
| Genome coverage (x) | 286.14 | 149.58 |
| Cytosine coverage (%) | 98.816 | 97.688 |
| Overall methylation level (%) | 0.523 | 0.564 |
| mC sites | 462 | 459 |
| mCG sites | 27 | 19 |
| mCHG sites | 19 | 24 |
| mCHH sites | 416 | 416 |

**Table 4.** Shared mCGs, mCHGs and mCHHs. The methylated sites identified in both replicate experiments are listed, including 5 mCGs, 2 mCHGs and 44 mCHHs, by genomic location, coordinate, total read number and methylation level. Many of these robust mCs show methylation levels above 10% in both experiments. Amongst the mCHHs, 20 are found in close proximity pairs (shaded boxes).

| Chromosome /<br>Contig | Genomic<br>Coordinate | First-round<br>Total Reads | Methyl Level (%) | Second-round<br>Total Reads | Methyl Level (%) |
| --- | --- | --- | --- | --- | --- |
| CG |  |  |  |  |  |
| 1 | 3783605 | 46 | 36.96 | 9 | 44.44 |
| 2 | 1836074 | 12 | 58.33 | 4 | 75.00 |
| 4 | 4598084 | 38 | 13.16 | 10 | 30.00 |
| MT | 28028 | 14 | 42.86 | 10 | 30.00 |
| Unmapped 13 | 3627 | 73 | 15.07 | 5 | 100.00 |
| CHG |  |  |  |  |  |
| 2 | 7430589 | 72 | 18.06 | 17 | 35.29 |
| 5 | 1898452 | 89 | 8.99 | 20 | 25.00 |
| CHH |  |  |  |  |  |
| 1 | 3379 | 195 | 5.13 | 55 | 10.91 |
| 1 | 9597 | 196 | 6.12 | 25 | 20.00 |
| 1 | 49448 | 171 | 5.85 | 22 | 18.18 |
| 1 | 1633944 | 134 | 21.64 | 53 | 15.09 |
| 1 | 1633981 | 150 | 15.33 | 65 | 23.08 |
| 1 | 2207539 | 74 | 40.54 | 17 | 23.53 |
| 1 | 3659412 | 65 | 16.92 | 24 | 37.50 |
| 1 | 4540349 | 28 | 17.86 | 17 | 29.41 |
| 2 | 493889 | 68 | 20.59 | 12 | 33.33 |
| 2 | 8101959 | 85 | 23.53 | 23 | 26.09 |
| 2 | 8102002 | 97 | 28.87 | 20 | 20.00 |
| 2 | 8269415 | 44 | 18.18 | 15 | 33.33 |
| 3 | 695099 | 62 | 14.52 | 17 | 23.53 |
| 3 | 799417 | 34 | 20.59 | 6 | 100.00 |
| 3 | 799419 | 54 | 33.33 | 22 | 31.82 |
| 3 | 1474373 | 90 | 12.22 | 22 | 27.27 |
| 3 | 3607695 | 109 | 8.26 | 45 | 24.44 |
| 3 | 4545189 | 53 | 11.32 | 17 | 23.53 |
| 3 | 5720793 | 74 | 29.73 | 16 | 37.50 |
| 3 | 5720812 | 84 | 11.90 | 15 | 26.67 |
| 3 | 5920588 | 97 | 15.46 | 23 | 17.39 |
| 3 | 5920623 | 82 | 18.29 | 20 | 20.00 |
| 3 | 6220024 | 21 | 19.05 | 10 | 30.00 |
| 4 | 76910 | 104 | 16.35 | 39 | 12.82 |
| 4 | 76914 | 105 | 20.95 | 41 | 12.20 |
| 4 | 76958 | 139 | 28.06 | 31 | 25.81 |
| 4 | 76962 | 138 | 20.29 | 34 | 14.71 |
| 4 | 2524803 | 112 | 32.14 | 46 | 28.26 |
| 4 | 2524824 | 117 | 31.62 | 45 | 24.44 |
| 4 | 4264271 | 33 | 27.27 | 5 | 60.00 |
| 4 | 4598089 | 36 | 13.89 | 12 | 33.33 |
| 4 | 5418388 | 90 | 10.00 | 23 | 21.74 |
| 4 | 5418407 | 65 | 10.77 | 6 | 50.00 |
| 5 | 1078684 | 86 | 12.79 | 16 | 31.25 |
| 5 | 2205539 | 23 | 26.09 | 7 | 57.14 |
| 5 | 4416786 | 311 | 15.11 | 425 | 11.29 |
| 5 | 4652725 | 47 | 21.28 | 10 | 30.00 |
| 6 | 617155 | 65 | 12.31 | 22 | 18.18 |
| 6 | 617176 | 57 | 15.79 | 24 | 45.83 |
| 6 | 1768503 | 303 | 9.24 | 465 | 4.52 |
| MT | 15196 | 16 | 100.00 | 5 | 100.00 |
| MT | 49661 | 9 | 44.44 | 23 | 21.74 |
| Unmapped 5 | 1301 | 113 | 14.16 | 22 | 27.27 |
| Unmapped 24 | 4083 | 73 | 13.70 | 8 | 75.00 |

**Figure 5.**
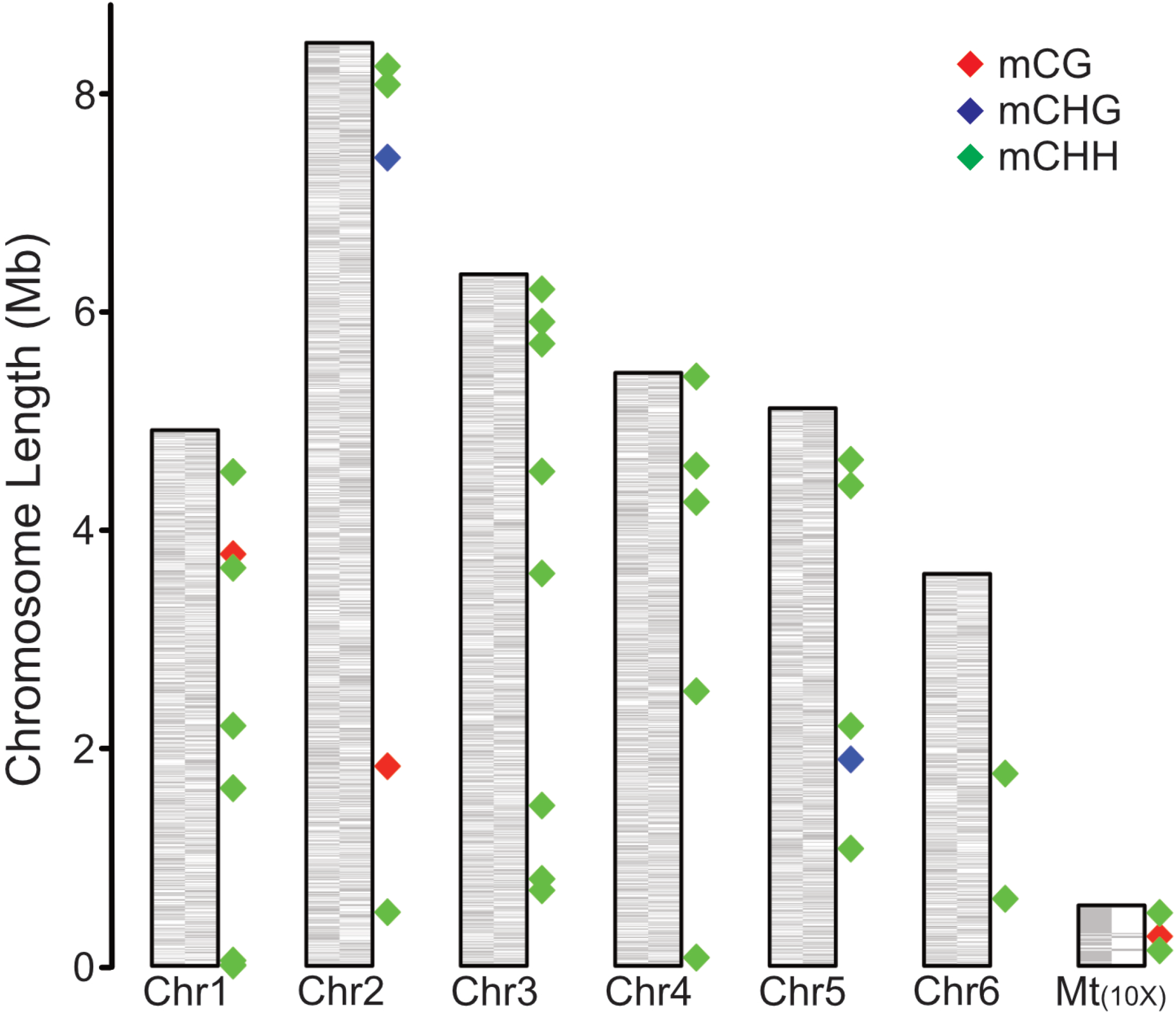
Genomic location of robust methylated sites. The six nuclear chromosomes and mitochondrial genome (10x enlarged) are depicted with plus strand (left side) and minus (right side) genes indicated (grey boxes). Diamonds depict the location of the robust CG (red), CHG (blue), and CHH (green) methylated sites detected in the replicate experiments. These methylated sites are distributed throughout the genome and occur predominantly at CHH sequences.

### Robust mC genomic features

Of the 51 robust mCs, 11 are located directly in annotated genes (Table S3). Gene Ontology (GO) analysis of these 11 mCs revealed enrichment in genes involved in the generation of precursor metabolites and energy (GO:0006091), driven by the *nad1* (*DDB_GO294018*), *nad11 (DDB_GO294052*), and *atp1 (DDB_GO0294012*) genes. Contrary to an earlier study [20], no evidence of DNA methylation at the *guaB* gene could be detected in our analysis. The nearest mC to guaB is over 1.5million base pairs away, suggesting that this gene is not a target for methylation in late development. Intriguingly, a single methylated site is detected in the DIRS1 retrotransposon on chromosome 1 (Table S3), supporting the previous observation of methylation in this class of retrotransposable element [20].

Examining the genomic neighborhoods around the 51 robust mCs, there are a total of 98 genes located within 1kb either upstream or downstream of individual mCs (Table S4). Amongst these genes, no single GO term is enriched and a wide range of protein classes are represented, including enzymes potentially involved in metabolic pathways (hydrolases, oxireductases and transferases) and nucleic acid binding proteins (Fig. 6). Given that our analysis was performed on the *Dictyostelium* cells at 18-24 hours post-starvation, it is perhaps not surprising to also find methylation proximal to, or within, genes associated with developmental-related processes such as slug and sorocarp stalk cell differentiation and morphogenesis (*proA/DDB_G0287125*, *stlB/DDB_G0290853 and DDB_G0277555*) and pheromone response (*DDB_G0284821*) (Table S3 and S4). A number of retrotransposable elements are also located within 1kb of a mC, including six from the DIRS1 family and three from the TRE family (Table S4), supporting a possible role for DNA methylation in the silencing of retrotransposons, as has been previously reported [18]. In addition, six different tRNA-encoding genes (*DDB_G0294885, DDB_G0295189, DDB_G0294641, DDB_G0294643, DDB_G0294645, and DDB_G0294657*) are located close to mCs (Table S4).

**Figure 6.**
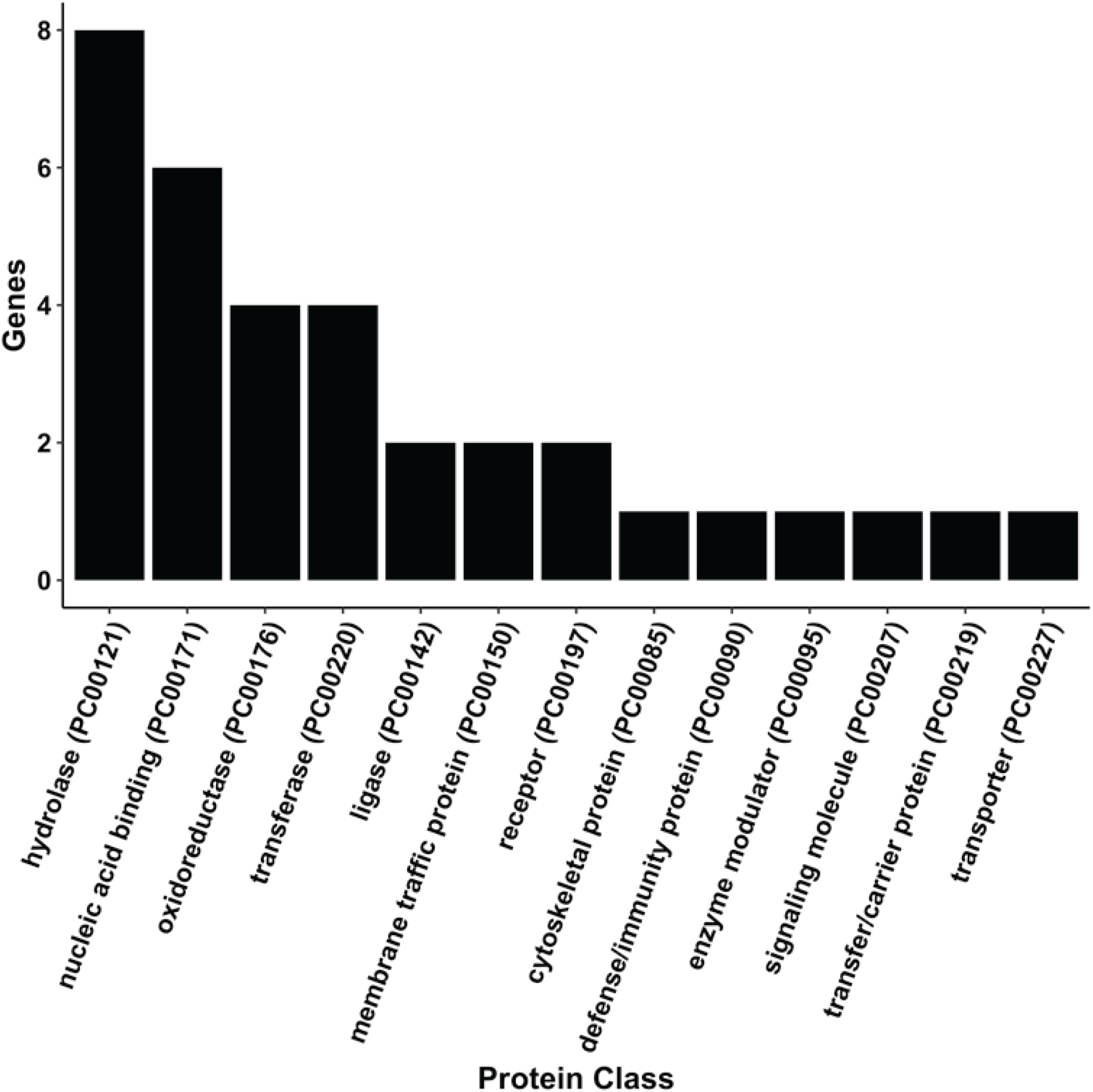
Protein classes of genes within 1kb of a robust methylated site. The frequency of different GO protein classes for the annotated genes located within 1kb of a robust methylated site are depicted. Methylated sites were most often associated with genes encoding for hydrolases (PC00121) and nucleic acid binding (PC00171).

## Conclusions

The paucity of studies investigating DNA methylation in *D. discoideum* has prohibited our detailed understanding of the evolution and function of eukaryotic DNA methylation with reference to D. *discoideum’s* unique phylogenetic position. That is, *D. discoideum* diverged soon after the plant-animal split and is therefore closely related to two kingdoms with largely present or absent DNA methylation systems (i.e., plants and fungi, respectively) [17] (Fig. 1). In our current study utilizing deep sequencing, we demonstrate that *D. discoideum* does possess a rudimentary DNA methylation system, as evidenced by overall very low, but reproducible levels of methylation in the genome. Our findings support the earlier observation that the genome of *D. discoideum* is largely devoid of methylation [19], but a few sites show robust, replicable patterns of methylation. Amongst these sites are mCs in, or in close proximity to, annotated transposable elements, tRNA genes and genes encoding for proteins with metabolic and developmental pathways [20]. Combining our new data with the knowledge that there is a functional DNMT2-homolog (DNMA) with potentially broad substrate recognition in *D. discoideum*, opens up the exciting possibility of further dissecting this evolutionarily conserved epigenetic system in future studies.

## Methods

### DNA Sources

To investigate the potential for DNA methylation sites in *D. discoideum*, AX4 strain cells were selected for whole genome analysis. *D. discoideum* AX4s have near-complete chromosome level genome assembly with genes functionally validated or predicted, making it the ideal and only current candidate for whole genome analysis. *D. discoideum* AX4s were grown in HL5 liquid cultures (dictybase.org for recipe) until near saturation. Once sufficient cell density was reached, 30mL of the culture was transferred to 50mL conical tubes. The cells were centrifuged for 5 minutes at 1000RPM, room temperature, the media was aspirated from the tube, and the cells were resuspended in 1.5mL 1x Starvation Buffer (dictybase.org). The resuspended cells were transferred to nitrocellulose membrane pads with 0.45μm pores pre-wetted with Starvation Buffer, and allowed to develop over the course of 18 to 24 hours. After this time, the cells were scraped from the pads, and gDNA was extracted with a Genesee Scientific ZR Genomic DNA Tissue MiniPrep extraction kit, utilizing the Solid Tissue protocol. After extraction, bisulfite sequencing was conducted to determine potential sites of methylation in the genome.

### Sequencing of Bisulfite Converted DNA Libraries

Library construction, bisulfiite conversion and sequencing were performed at the Beijing Genomics Institute. Briefly, DNA was fragmented into 100-300 bp fragments by sonication (Covaris S-2, Woburn USA). The fragmentation parameters were: Duty cycle 10%; Intensity: 5; Cycles/burst: 200; Cycles: 16; Total fragmentation time: 960 sec. Fragmentation was confirmed using a 2100 Bioanalyzer (Aligent Technologies, Santa Clara, USA). Fragments were end repaired (Illumina) as recommended by the manufacturer. Repaired fragments were ligated with methylated sequencing adaptors using a paired end adaptor oligo kit and oligo mix 5 (Illumina). Ligated fragments were selected by gel electrophoresis and fragments of size 360 bp extracted using a QIAquick gel extraction kit (Qiagen).

Size-selected fragments were bisulfite treated using an EZ-DNA methylation kit (Zymo Research, Irvine USA) and enriched using a MethyMiner methylated DNA enrichment kit (Invitrogen). It should be noted that this kit uses the DNA binding domain from human methyl-binding domain 2 protein to enrich for methylated DNA and therefore, when compared to experimental approaches that do not use this step, it will likely introduce selection for methylated fragments. Libraries were amplified using T4 polymerase (Enzymatics), and sequenced on Illumina’s HiSeq PE 150 platform.

### Sequence Analysis and Mapping DNA Methylation

Data were filtered to remove adaptor sequences, duplicate sequences, contamination and low quality reads using BGI software. We mapped our reads onto the *Dictyostelium discoideum* genome assembly 1.0 [17] using BSMAP version 2.6 [26] with seed size 12 and maximum allowed mismatches 5. Similarly, we mapped our reads onto the complete lambda phage genome (GenBank: J02459.1). 72.53% of our reads mapped onto the *Dictyostelium* genome. We considered only reads that mapped uniquely, and bases within reads that had a quality score of 20 or more, and that were next to 3 matches with quality scores of at least 15 [27]. From these data, we determined the number of converted and unconverted reads at each cytosine position in the *Dictyostelium* and lambda genome assemblies, accounting for the fact that each read comes from a bisulfite reaction on one strand or the other.

To estimate the overall rate of bisulfite conversion in non-methylated bases in our experiments, we used the C to T conversion rate in the lambda phage DNA, in which all cytosines should have been converted. We found that 99.44% of cytosines were converted in the lambda DNA, indicating that our false negative rate is less than 0.6%.

The average methylation level by chromosome was determined by the ratio of the number of reads supporting methylation to the number of reads covering a particular cytosine site:

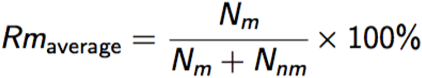

Where N_m_ represents the number of methyl-C (non-converted) reads and N_nm_ represents the number of non-methylated (converted) reads.

To identify individual cytosines that were significantly methylated in the *Dictyostelium* genome, we compared the number of converted and non-converted reads at each site. We used only sites that had coverage of 4 or more and no more than 1000 reads. We asked how likely these counts were under a binomial test where the probability of success is one minus the conversion rate, and corrected this probability value for multiple testing [28], as previously described [24, 25, 29]. From this, were able to determine the methylated sites in the genome and the level of methylation at individual 5-methylcytosines.

### Gene Ontology Enrichment and Protein Class Mapping

To determine what functional categories are overrepresented among methylated sites of genes either directly methylated or within 1kb of a methylated site, a list of genes fitting these criteria were identified using BEDTools, version 2.25 [30]. The list of genes was then inputted to AmiGO2, version 2.4.24 [31] using the PANTHER, version 11.1 [32], overrepresentation test (release 20160715). Similarly, to summarize the protein classes of genes within 1kb of a methylated site, PANTHER’s functional classification function was used.

## Supporting Information

**Supplementary Table 1. Cytosine coverage by chromosome**

The values represent the average coverage at cytosine bases on each chromosome and the mitochondrial DNA (chrMT). The profile for cytosines in different sequence contexts (CG, CHG and CHH, where H = non-G base) is shown.

**Supplementary Table 2. Robust mCHH proximity pairs GO classification**

The chromosomal location and identity of different GO protein classes for the annotated genes located within 1kb of a robust methylated CHH proximity pair are listed.

**Supplementary Table 3. 11 robust mCs are located directly in genes**

The chromosomal location, sequence context and identity for the annotated genes are shown.

**Supplementary Table 4. Genes within 1kb window of a robust mC**

The chromosomal location, sequence context and identity for the annotated genes within 1kb of a robust methylated cytosine are listed.

## Availability of Supporting Data

The data set supporting the results of this article is available at: Will be publicly released at publication

## Abbreviations

DNMT = DNA methyltransferase

GO = Gene ontology

## Competing interests

The authors report no competing interests.

## Authors’ contributions

JLS conceived of the study, participated in the design of the study and helped to draft the manuscript. JSD performed the gDNA preparation and drafted the manuscript. JMD carried out sequence and bioinformatic analysis. DAL conceived of the study, participated in the design of the study and helped to draft the manuscript. RAD conceived of the study, participated in its design, established the genomics pipeline, coordinated and drafted the manuscript. All authors read and approved the final manuscript.

## Acknowledgements

This work was funded in part by a National Institutes of Health (GM110571) grant to RAD and JMD.

